# The songbird lateral habenula projects to dopaminergic midbrain and is important for normal vocal development

**DOI:** 10.1101/2023.06.21.545765

**Authors:** Andrea Roeser, Han Kheng Teoh, Ruidong Chen, Itai Cohen, Jesse Goldberg

## Abstract

Mistakes in performance feel disappointing, suggesting that brain pathways for aversive feedback may play a role in motor learning. Here we tested if the lateral habenula (LHb), an evolutionarily conserved part of the limbic system known in mammals to relay aversive feedback from ventral pallidum (VP) to ventral tegmental area (VTA) dopamine neurons, is involved in birdsong learning and production. By combining viral tract tracing and functional circuit mapping, we discovered that songbird LHb links VP and an auditory cortical area to singing-related DA neurons that signal song errors. As in mammals, VP stimulation activated LHb activity and LHb stimulation suppressed DA firing. To test this pathway’s role in learning we lesioned the LHb in juvenile zebra finches and recorded their songs in adulthood. Birds with the LHb lesioned as juveniles produced highly unusual vocalizations as adults, including prolonged high-pitch notes and species-atypical trills. These findings identify a songbird VP-LHb-VTA pathway with similar functional connectivity as mammals, expand the known territories of vocal learning circuits, and demonstrate that limbic circuits associated with disappointing outcomes are important for motor performance learning.

## Introduction

Dopamine (DA)-basal ganglia (BG) circuits are critical for motor control and learning. During reward seeking, DA activity is activated by surprisingly good reward outcomes and is decreased by disappointingly bad ones (Schultz et al., 1997). These ‘reward prediction error’ (RPE) signals mediate reinforcement learning by regulating plasticity in BG motor circuits (Costa, 2007; Wickens et al., 2003). Yet it remains unclear how mechanisms of reinforcement learning identified in reward-seeking animals relate to natural behaviors with no primary rewards at stake.

Songbirds offer a unique opportunity to study internally-guided motor learning. Song is learned not for external reinforcement but instead by matching vocal performance to the memory of a tutor song (Marler, 1997, 1970). Song learning also requires a DA-BG circuit that is part of a tractable song system, which includes a pathway from the ventral tegmental area (VTA) to the song-specialized striatopallidal nucleus, Area X (Chen and Goldberg, 2020; Person et al., 2008). Area X-projecting VTA neurons are dopaminergic (Person et al., 2008; Reiner et al., 2004), regulate synaptic plasticity in Area X (Ding and Perkel, 2004), are necessary for normal song learning (Hoffmann et al., 2016; Saravanan et al., 2019), and exhibit RPE-like performance error signals during singing that can reinforce song variations. Specifically, in experimental paradigms where specific song syllables are negatively reinforced with distorted auditory feedback (DAF), distorted syllable renditions evoke phasic suppression of DA activity that extinguish those variations, and undistorted renditions evoke activation of DA activity and syllable reinforcement (Duffy et al., 2022; Gadagkar et al., 2016; Hisey et al., 2018; Xiao et al., 2018).

Given the conserved roles of DA signals in birdsong and reinforcement learning (Chen and Goldberg, 2020), inputs to mammalian VTA provide a roadmap for the discovery of pathways in songbirds that may contribute to dopaminergic song evaluation. Consistent with deep homology in vertebrate brain architecture (Reiner, 2009; Swanson, 2000), we and others recently discovered three conserved inputs to VTA that are important for song learning: the ventral pallidum (VP), the subthalamic nucleus (STN), and an auditory error-associated cortical area (Bottjer et al., 2000; Chen et al., 2019; Das and Goldberg, 2022; Gale and Perkel, 2010; Kearney et al., 2019; Mandelblat-Cerf et al., 2014). First, VP lesions impair song learning, optogenetic manipulation of the VP-VTA pathway reinforces song syllable variations, and VTA-projecting VP neurons exhibit DAF-associated auditory error signals during singing, including temporally precise information about predicted song quality (Chen et al., 2019; Kearney et al., 2019). Second, the STN receives input from VP and projects to VTA, and STN neurons can exhibit singing-related activity, including response to DAF-based song errors (Das and Goldberg, 2022). Finally, a high-order auditory cortical area, the ventral intermediate arcopallium (AIV), is important for song learning and also sends singing-related DAF auditory error signals to VTA that can influence song learning (Bottjer et al., 2000; Kearney et al., 2019; Mandelblat-Cerf et al., 2014). Though cortical (pallial) songbird regions have complex and unclear analogies to the mammalian cortex, several high-order cortical regions, including cingulate regions activated by prediction errors, project to VTA (Elston and Bilkey, 2017). These similarities in songbird and mammal input pathways to VTA led us to hypothesize that songbird lateral habenula (LHb) may, as in mammals, also project to VTA and contribute to learning-related DA signals (Fig. 1D).

**Figure 1.**
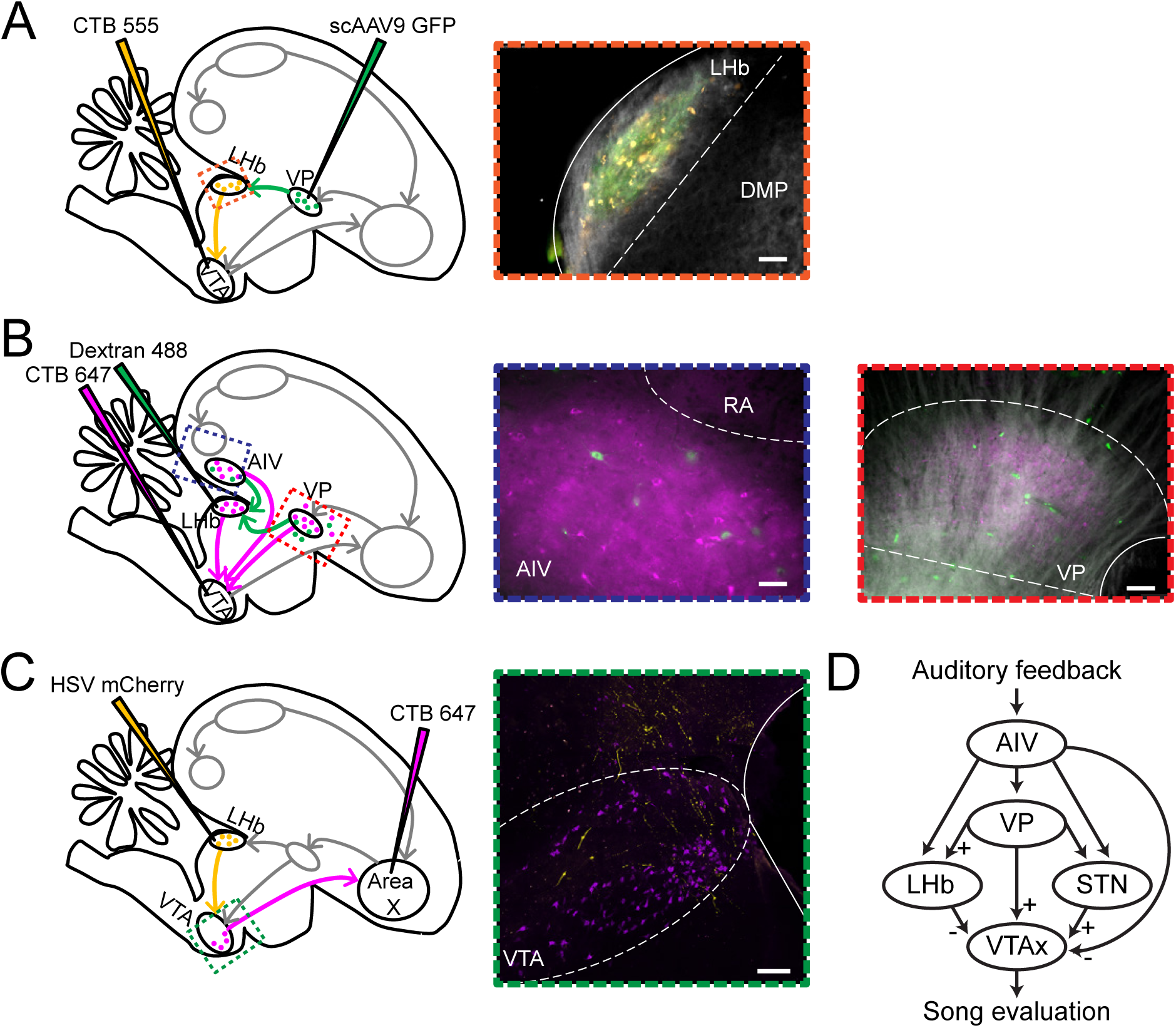
LHb receives inputs from VP and AIV and projects to VTA. A: Injection of viral tracer scAAV9-CBh-GFP into VP, and retrograde tracer cholera toxin subunit B (CTB) 555 into VTA resulted in co-localization of VP axons (green) and VTA projecting neurons (yellow) in LHb. Scale bar: 50 um. B: LHb- and VTA-projecting neurons (green and magenta, respectively) were identified in both VP and AIV following injection of retrograde tracer Dextran 488 into LHb and CTB 647 into VTA. Note LHB- and VTA-projecting neurons spatially intermingled in both VP and AIV but rarely co-labeled. Scale bar: 50 um (center), 75 um (right). C: LHb axons (yellow) innervated the area of Area X projecting VTA neurons (magenta) following injection of CTB 647 into Area X and viral tracer HSV-mCherry into LHb. Scale bar: 75 um. All histological images are sagittal slices with dorsal upwards and anterior to the right. D: A summary schematic of input pathways to songbird and mammalian VTA. The excitatory (+) or inhibitory (-) nature of the functional connections were established in mammals and, more recently, in songbirds (see also Fig. 2 of this paper).

The LHb-VTA projection is conserved from fish to humans (Boulos et al., 2017; Namboodiri et al., 2016; Ullsperger and von Cramon, 2003). Across model systems and studies, LHb neurons are activated by disappointing reward outcomes and aversive stimuli (Andalman et al., 2019; Matsumoto and Hikosaka, 2009). LHb lesions impair DA reward processing, especially reward omission-associated signals (Tian and Uchida, 2015). Given the importance of the LHb in DA signaling, and the importance of DA signaling for song learning, we investigated LHb function in songbirds for the first time.

Here we combine neural tracing, functional mapping, and lesion studies in juvenile and adult birds to test LHb’s connectivity and its role in song learning. We find that AIV and VP send projections to LHb, and that LHb microstimulation evokes phasic suppression in Area X projecting VTA neurons. Finally, consistent with a role for LHb in song learning, LHb lesions in juvenile birds cause abnormal vocalizations, but lesions in adults had no effect on the production of already-learned songs.

## Methods

### Subjects

Subjects were 59 male zebra finches (at least 39 days post hatch, dph). Animal care and experiments were carried out in accordance with NIH guidelines and were approved by the Cornell Institutional Animal Care and Use Committee.

### Surgery and Histology

For every surgery, birds were anesthetized with isoflurane. Anatomical coordinates are relayed as the distance, antero-posterior (A) and mediolateral (L), from the bifurcation of the sinus vein (lambda), and dorsoventral (V) from the pial surface.

For tracing experiments, 14 birds that were 90 dph or older, were used to determine the connectivity between LHb and the song system. To test if VP axons co-mingle with VTA projecting LHb neurons (7 birds, 9 hemispheres), 40 nL of self-complementary adeno-associated virus carrying green fluorescent protein (scAAV9-CBh-GFP, UNC Vector Core) was injected into VP (20 degree head angle; 4.9A, 1.3L, 3.8V) and 40 nL of fluorescently labeled cholera toxin subunit B (CTB, Molecular Probes) was injected into VTA (40 degree head angle; 3.4A, 0.6L, 6.5V). To determine if the projections from AIV and VP to both VTA and LHb are distinct or have collaterals (4 birds, 6 hemispheres), 40 nL of CTB647 was injected into VTA and 40 nL of CTB488 or Dextran 488 (10,000MW, ThermoFisher) into LHb (75 degree head angle; 0.3A, 0.6L, 3.3V). To test if LHb axons localized with Area X projecting VTA neurons (2 birds, 3 hemispheres), 30 nL of HSV-mCherry (MGH Viral Core) was injected into LHb, and 50 nL of CTB647 into Area X (20 degree head angle, 5.6A, 1.5L, 2.8V). Injections into VP, VTA, and Area X were made with a Nanoject II (Drummond Scientific, Broomall, PA), and injections into LHb were accomplished with a Picospritzer II (Parker Hannifin Co., Hollis, NH) and ejection micropipette (Carbostar-3, Kation Scientific).

Functional mapping experiments were conducted to determine the connectivity between three different brain regions: VP and LHb, and LHb and VTA. Recordings from LHb and VTA neurons were made using a carbon fiber electrode (Carbostar-1, 1 MOhm, Kation Scientific), while birds were anesthetized. In the first experiment, bipolar stimulation electrodes were implanted into VP (n = 3 birds) and in 2/3 birds, an additional stimulation electrode was implanted into VTA (same coordinates used for tracing experiments). To find LHb coordinates, the posterior boundary of DLM was found using the distinctive spontaneous firing rate found within that brain area. LHb was then recorded from roughly 300 um posterior to the posterior edge of DLM, 0.6L, and 3.8V, at a head angle of 55 degrees. All neurons’ responses to VP stimulation were recorded (n = 8 neurons), and in the 2 birds with VTA stimulation electrodes, all neurons (n = 6 neurons) were tested for antidromic responses that were also confirmed with collisions. In the second experiment, bipolar stimulation electrodes were implanted into both LHb and Area X (same coordinates used for tracing experiments; n = 5 birds). All VTA neuronal responses to LHb stimulation were recorded (same coordinates used for tracing experiments; n = 19 neurons), and Area X-projecting VTA neurons were confirmed by antidromic response and further collision testing. At the end of each experiment, to verify location of stimulation and recording sites, small lesions were made by passing 30 uA of current for 60 s at each bipolar electrode stimulation site, as well as one neuronal recording site for each bird.

Prior to all LHb lesion experiments, birds used for juvenile lesions were hatched and housed in a colony of conspecifics with continuous access to a tutor. Juveniles were separated from the colony (37 - 43 dph) and housed in acoustically isolated recording boxes for the remainder of the experiment. Vocalizations were recorded for the duration of the experiment using Sound Analysis Pro (SAP) software in sound isolation boxes (Tchernichovski et al., 2000). Once a full day of vocal babbling was recorded, birds were considered ready for surgery. Birds used for adult lesions were also housed in a colony of conspecifics prior to the start of the experiment, then acoustically isolated in recording boxes for the duration of the experiment. At least five days of singing was recorded before lesion surgery (>150 dph; n = 5 adult birds).

To ensure the accuracy and extent of the lesions, coordinates and microlesion locations from the functional mapping birds described above, and 3 additional adults (6 hemispheres) were used to test and calibrate the location and lesion amount due to varying current intensities. The lesion surgery began by mapping the posterior edge of DLM (as described for functional mapping experiments) using a carbon fiber electrode. Then in 6 juvenile birds (43-53 dph), roughly 150 nL of 2% N-methyl-DL-aspartic acid (NMDA; Sigma, St Louis, MO) was injected bilaterally into the LHb using a Picospritzer II and ejection micropipette. A further 7 juvenile birds were injected with saline (for sham birds). In the remaining juvenile and adult birds, a carbon fiber electrode was used to electrolytically lesion LHb by bilaterally passing ∼40 uA of current for ∼65 s in 2-3 locations (n = 11 juvenile birds 39-52 dph; n = 5 adult birds >150 dph), or the electrode was inserted into LHb but no current was passed (sham birds, n = 2 juvenile birds, 41-42 dph). Lesions were targeted to the LHb, but we cannot rule out the possibility that medial habenula was partially lesioned in some hemispheres. Lesions were visually confirmed by observing a pronounced absence of cells and tissue damage compared to sham birds using anti-NeuN (neuronal nuclear stain; n = 12/17 juvenile; n = 9/9 sham; n = 3/5 adult birds) and/or bilateral injections of CTB 647 into VTA to test for any remaining VTA projecting LHb neurons (n = 15/17 juvenile; n = 2/9 sham; n = 5/5 adult birds). Because of the months between lesion surgery and histological verification (to allow for vocal development), precise lesion size quantification was not possible. However tissue damage and cell loss was confirmed for every bird in the lesion dataset.

All birds were perfused with a 4% paraformaldehyde solution and brains were sectioned into 100 um sagittal slices for imaging. All imaging was acquired using a Leica DM4000 B microscope, except for imaging LHb axons in VTA, which was taken using a Zeiss LSM 710 confocal microscope (Fig. 1C).

### Functional Mapping and Analysis

In both LHb and VTA, neurons were deemed VTA-projecting LHb (LHb^VTA^) or Area X-projecting VTA (VTA_X_) neurons based on antidromic stimulation and collision testing (200 us pulses, 100-300 uA). Neurons not tested for antidromic responses (n = 2 LHb neurons), or those that did not respond to VTA or Area X stimulation were classified as LHb_other_ or VTA_other_ neurons, however, there is the possibility that some of these neurons project to areas outside of the area of stimulation and in fact are LHb_VTA_/VTA_X_ neurons. VP stimulation of LHb neurons was a single stimulation (200 us, 200-300 uA current amplitude) delivered roughly every 1.5-2 s (Fig. 2 A-G). LHb stimulation of VTA neurons was a burst of three 200 us pulses with a 3 ms inter-pulse-interval delivered roughly every 2 seconds (200-300 uA; Fig. 2 H-Q). Spike duration of VTA neurons was calculated by finding the interval between spike onset and offset (Fig. 2J).

**Figure 2.**
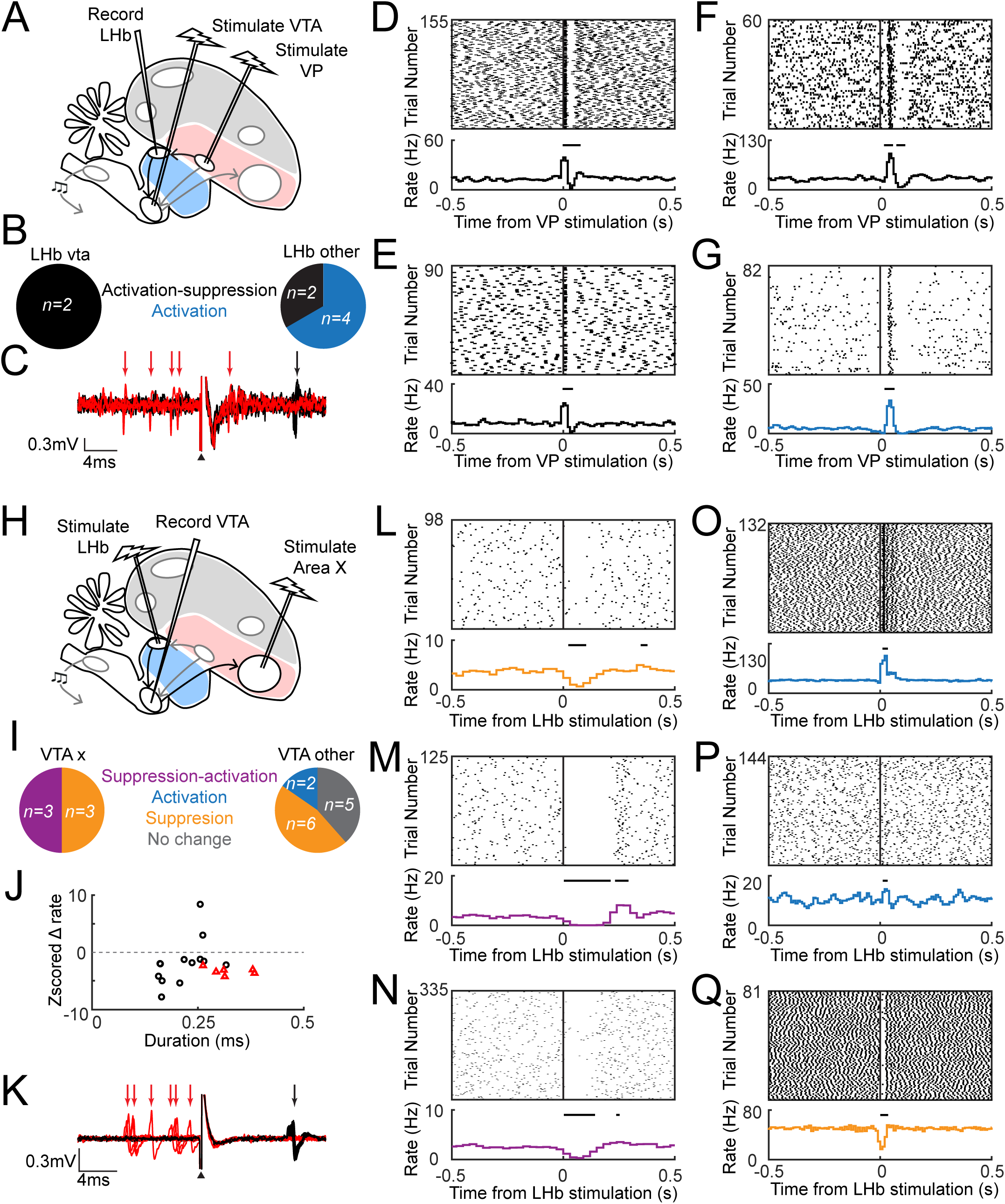
VP activates LHb, which suppresses Area X-projecting VTA DA neurons. A: Brain schematic of the experimental approach to test how VTA-projecting LHb neurons (LHb_VTA_) respond to VP stimulation. B: Summary of LHb_VTA_ (left) and LHb_other_ (right) responses. C: antidromic identification of an LHb_VTA_ neuron. Collisions (red) and antidromic (black) spikes are indicated by arrows, and the stimulation artifact by the black triangle. D: example LHb_VTA_ neuron (same as in C). Raster plot (top) and rate histogram (bottom) aligned to a single VP stimulation pulse. Horizontal bars show a significant response (p < 0.05, Z test; Methods). E: data plotted as in D for a second LHb_VTA_ neuron. F and G: data plotted as in D for two LHb_other_ neurons. H: brain schematic of the experimental procedure to record the responses of Area X projecting VTA neurons (VTA_X_) to LHb stimulation. I: summary of VTA_X_ (left) and VTA_other_ (right) responses to burst LHb stimulation. J: scatterplot showing the Z-scored firing rate change within a 100 ms window from burst LHb stimulation against the average spontaneous spike width duration (VTA_X_ neurons: red triangles; VTA_other_ neurons: black circles). K: data plotted as in C for an antidromically identified VTA_X_ neuron. L: example VTA_X_ neuron (same as in K). Raster plot (top) and rate histogram (bottom) aligned to a triple LHb stimulation (interpulse interval, 3 ms). Horizontal bars show a significant response (p < 0.05, Z test). M and N: data plotted as in L for two additional VTA_X_ neurons. O-Q: data plotted as in L for three VTA_other_ neurons.

Responses of LHb and VTA neurons to stimulation were analyzed offline using custom MATLAB code (Chen et al., 2019; Das and Goldberg, 2022). Raster plots and peri-stimulus rate histograms (PSTHs) aligned to VP and LHb stimulation events were generated to assess LHb and VTA neuronal responses, respectively. To calculate the PSTHs and test for significance, all neuronal activity within one second of stimulation onset was binned in a moving window (5 ms step size). Due to differences in spontaneous firing rates, two different bin sizes were used: 30 ms for VTA_X_ neurons, and 10 ms for VTA_other_ and LHb neurons (Chen et al., 2019). After stimulation onset, each bin was tested for a significant change against all bins found in the previous 1 second(z-test, p < 0.05). If at least 2 consecutive bins had a p < 0.05, that window was considered significant (Fig. 2) and the latency to response was defined by the onset of the first bin.

### Song Similarity and Abnormal Vocalizations Analysis

All adult and juvenile LHb lesion birds were presented with a female and sang female-directed song (adult: > 150 dph, n = 5 adult; juvenile: 91-107 dph, n = 17 juvenile birds). Female-directed song was elicited from 4 juvenile sham birds (91-107 dph); the remaining 5 sham birds were never presented with a female, so no female-directed song data was available.

Computing a song imitation score allowed us to assess 1) the learning of the tutor song by LHb lesion and sham birds (juvenile LHb lesion bird song from 90 dph; Fig 3H), and 2) similarity of adult LHb lesioned bird song 1-2 days prior to lesion vs. 60 days post-lesion (Fig. 3D, red) as compared to pre-lesioned song taken 1-2 days and 3-4 days prior to lesion (Fig. 5D, black center) and song recorded 60 days apart from 3 adult control males (Fig. 5D, black left). The SAP algorithm, previously described (Tchernichovski et al., 2000), was used to compute imitation scores through an automated MATLAB GUIincluding only syllables found in the motif. The imitation score (compared to the tutor or self) is the product of acoustic and sequence similarity scores. In short, the template song (tutor or pre-lesion song) was segmented by hand into motif syllables. The acoustic similarity score was computed by comparing each template syllable with the part of the the pupil or post-lesion adult bird’s song that best matched it. Sequence similarity score was derived by comparing the similarity of the pupil/post-lesion syllable sequence to that of the template motif (Fig. 3H and Fig. 5D).

**Figure 3.**
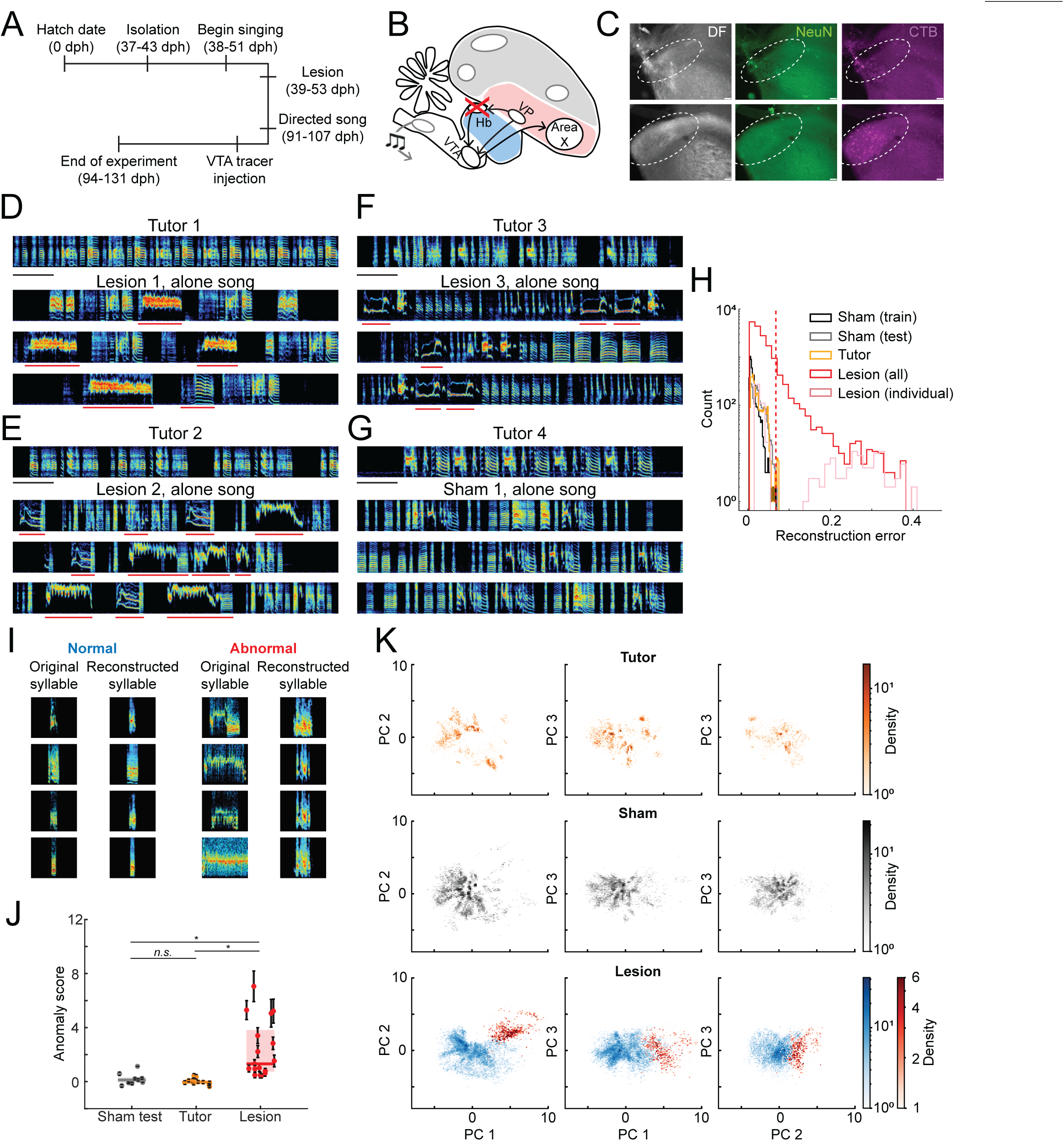
LHb lesions in juvenile birds cause atypical vocalizations in adults. A: experimental timeline. B: schematic showing the location of habenula in the diencephalon. C: lesions were confirmed using dark field (DF, left column), anti-NeuN (middle column), and CTB647 to label VTA-projecting LHb neurons (right column), LHb indicated by white circle. Top row: example of a completely lesioned hemisphere as evident by extensive tissue damage (small, bright dots in all three images), no neurons visible in anti-NeuN, and absence of LHb_VTA_ neurons. Bottom row: example of a sham bird with intact LHb at roughly the same mediolateral position as hemisphere in the top row. Note the cluster of LHb_VTA_ (magenta) absent in the fully lesioned condition. Scale bars: 75 um. D: spectrograms showing tutor song (top row), and adult undirected song of an LHb lesioned juvenile bird (bottom three rows). Spectrograms in D-G have frequency limits of 500-8500 Hz and are 4 seconds in length. Scale bar: 500 ms. E-F: data plotted as in D for two additional LHb lesioned juveniles. Red underlines indicate abnormal vocalizations G: data plotted as in D-F except for a sham bird. H: reconstruction error distributions for all datasets from one of the six replicates of a trained VAE. The red vertical line corresponds to the 99.9th percentile of sham bird (test set) vocalizations, which was used to define a boundary between normal and abnormal vocalizations (Methods). Sham birds (training set: black outline; test set: gray), LHb lesion birds (average: red; individual bird shown in D: pink), and tutor bird (yellow) vocalizations. I: example original and reconstructed spectrograms from the VAE of LHb lesioned birds with reconstruction error values less than the threshold (normal) and greater than the threshold (abnormal). J: average anomaly scores for each individual bird across 6 replicate VAEs (n = 9 sham birds; n = 12 tutor birds; n = 17 lesion birds; mean +/- STD shown in dot/vertical line). Lesion group is significantly different from the other two groups (sham and tutor; Kruskal-Wallis test with Bonferroni method multiple comparison: α = 0.05). K: two-dimensional projection of the PC space for the first three components. Top row corresponds to tutor vocalizations, middle row corresponds to all sham bird vocalizations (training and test set), and bottom row corresponds to LHb lesioned birds vocalizations (blue: normal syllables; red: abnormal syllables as determined by the threshold in H).

**Figure 4.**
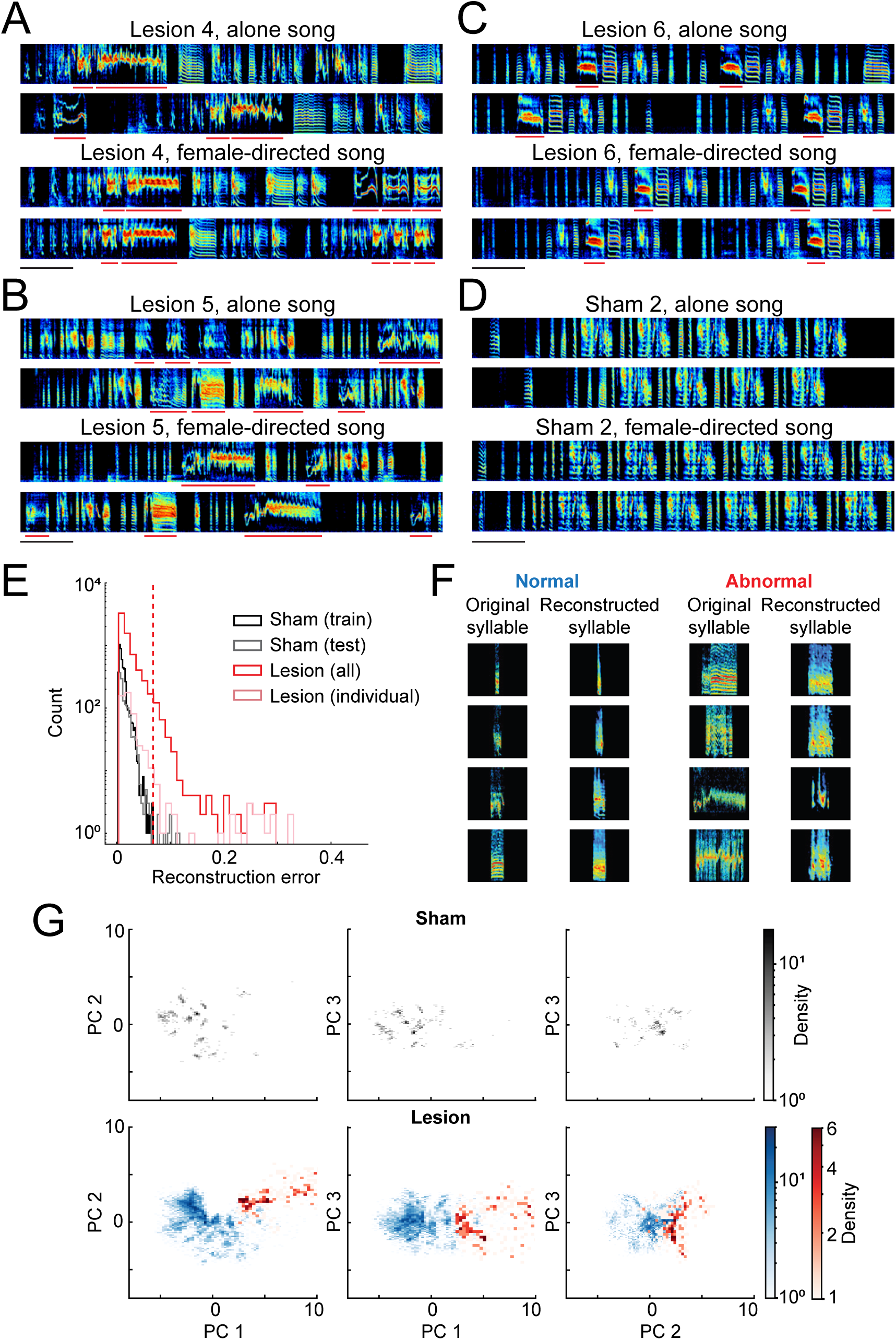
LHb-lesioned birds produce abnormal vocalizations during courtship song. A: spectrograms showing undirected song for a juvenile LHb lesion bird (top) and female-directed song for the same bird (bottom; bird not shown as an example in Fig 3). Scale bar: 500 ms. Red underlines indicate abnormal vocalizations. B-C: same as A for two additional LHb lesion birds, not shown as examples in Fig. 3. D: same as A-C except for a sham bird, not shown as an example in Fig. 3. E: reconstruction error distribution for sham bird (training set: black; test set, gray), and the LHb lesioned bird’s female-directed vocalizations (average: red; individual bird shown in A: pink). The red vertical line corresponds to the threshold set at 99.9th percentile of sham lesioned controls (test set) vocalizations. F: example original and reconstructed spectrograms from the VAE of LHb lesion birds with reconstruction error values less than the threshold (normal) and greater than the threshold (abnormal). G: two-dimensional projection of the PC space for the first three components. Top row corresponds to sham bird female-directed vocalizations (training and test sets), and bottom row corresponds to LHb lesion bird female-directed vocalizations (normal syllables: blue; abnormal syllables: red, as determined by the threshold in E).

**Figure 5.**
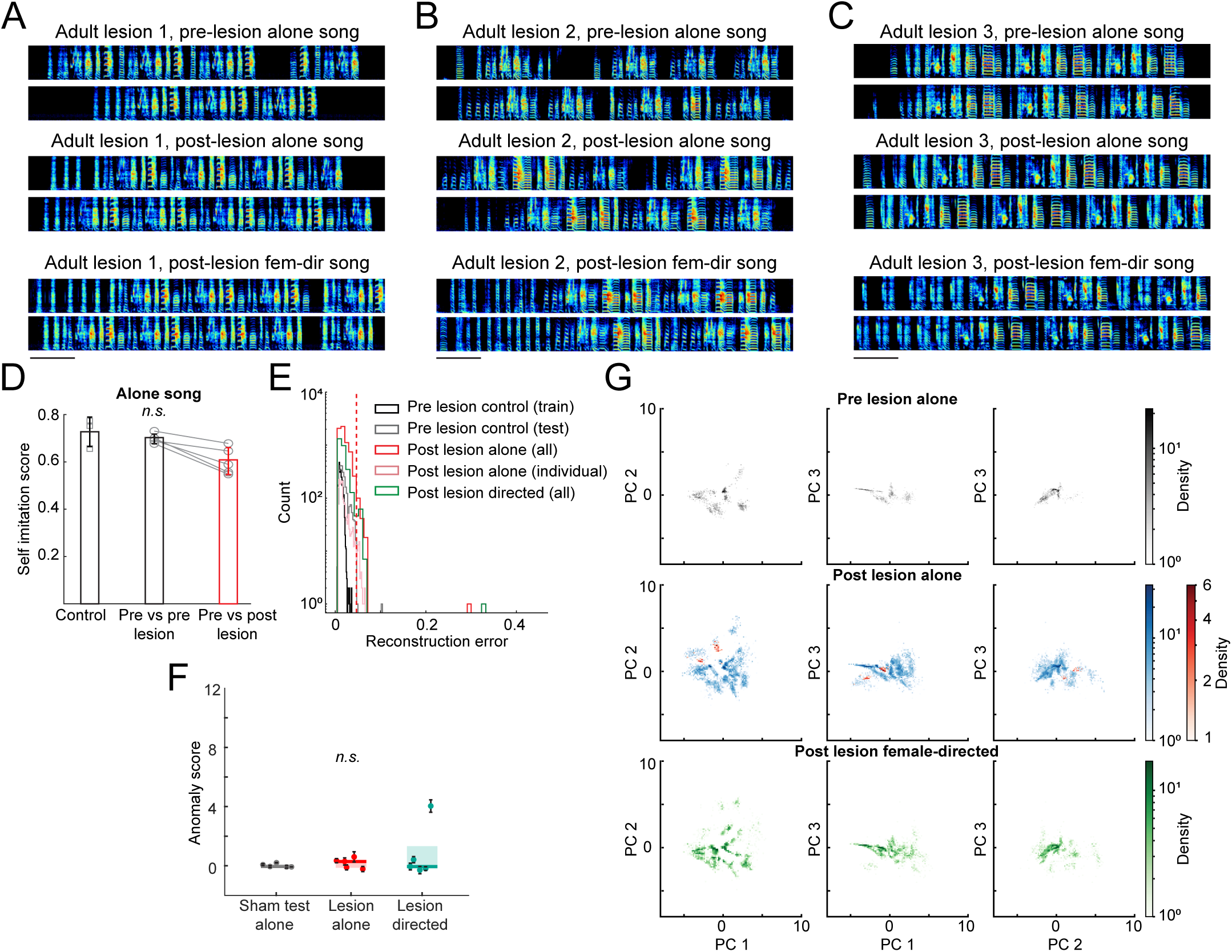
LHb lesions in adulthood do not impair vocalizations or the ability to produce female-directed song. A: spectrograms showing pre-lesion lone song for an adult LHb lesion bird (top), 60 days post-lesion lone song (middle), and post-lesion female-directed song for the same bird (bottom). Scale bar: 500 ms. B-C: same as A except for two additional example adult lesion birds. D: the self-imitation score was high for each bird, and there was no significant difference between any of the three groups (n = 3 birds control; n = 5 birds pre- vs. pre-lesion and pre- vs. post-lesion; p = 0.0625, two-sided, paired Wilcoxon sign rank test comparing pre- vs pre-lesion and pre- vs. post-lesion; p = 0.571, two-sided, unpaired Wilcoxon rank sum test comparing pre- vs. post-shamand pre- vs. pre-lesion; p = 0.0714, two-sided, unpaired Wilcoxon rank sum test comparing pre- vs. post-shamand pre- vs post-lesion). E: reconstruction error distribution for all vocalizations. The red vertical line corresponds to the threshold set at 99.9th percentile of pre-lesioned undirected song (test set), which is used to divide the LHb post-lesion undirected data into normal and abnormal vocalizations. LHb pre-lesion undirected (training set: black outline; test set: gray), LHb post-lesion undirected (average: red; individual bird shown in A: pink), and LHb post-lesion female-directed (green) vocalizations. F: average anomaly score for each individual bird across 6 replicate VAEs (n = 3 control; n = 5 lesion undirected; n = 5 lesion female-directed; mean +/- STD shown in dot/vertical line). Groups are not significantly different from each other (Kruskal-Wallis test, p = 0.9139). The outlier observed in the lesion directed group is due to determining the 99.9th percentile with a small sample size. The outlier (green) can be observed in E and Sup Fig 5_1A and C. G: two-dimensional projection of the PC space for the first three components. Top row corresponds to pre- lesion undirected vocalizations (test and training sets), middle row corresponds to post-lesion undirected vocalizations, and bottom corresponds to LHb post-lesion female-directed vocalizations.

### Segmentation and Spectrogramming

Using a custom-made MATLAB GUI, vocalizations were manually segmented for each bird: i) 2000 syllables from 5 consecutive days (90-94 dph) for juvenile LHb lesion and sham birds, ii) all tutor song from one day, iii) all female-directed song from one day from juvenile LHb lesion and sham birds, iv) 1500 syllables from 2 days pre and 1 day post adult lesion song, and v) all directed syllables for post adult lesion female-directed song. To create syllable spectrograms, log modulus of a syllable’s Short Time Fourier Transform (STFT) was computed using Hann windows of length 512 and overlap of 256. Sampling rate was 44100 kHz for all audio recording. The spectrogram’s frequency range was linearly spaced from 0.3 to 12 kHz. To ensure all syllable spectrograms have the same size, short syllables were zero-padded symmetrically to the size of the longest syllable in our segmentation. The resulting spectrograms were then rescaled to the intensity interval [0,1] and resized to 64 (frequency) by 256 (time) pixels.

## Training procedure of Variational AutoEncoders (VAEs)

To ensure a balanced training set for the VAE, we performed UMAP (McInnes et al., 2018), a non-linear dimensionality reduction technique on the syllable spectrograms for each bird, resulting in a two-dimensional projection. The syllables’ UMAP projections were subsequently clustered in an unsupervised manner via HDBscan (McInnes et al., 2017). This procedure yielded 4-7 clusters from each bird (two examples shown in Sup Fig 3_2 for birds in Figure 3). One hundred example spectrograms from each cluster were then randomly selected to form part of the training and test sets. Five sham birds’ syllable spectrograms were selected as our training set for the juvenile-lesion song analysis, and 3 pre-lesioned adult-lesion birds’ syllable spectrograms were used as our training set in adult song analysis. As some of the sham birds had LHb-lesioned siblings who learned from the same tutor, we ensured the training set had a proper ratio of birds with and without LHb-lesioned siblings in the test set. The remaining spectrograms were used as the test set. To ensure our result was robust against selection bias, we performed six-fold cross validation for both juvenile-lesion (Supplementary Table I, Fig 3 and 4, and Sup Fig 3_2) and adult lesion song analysis (Supplementary Table II, Fig 5, and Sup Fig 5_1) by permuting the sham birds across training and test sets.

The VAE network was implemented and trained in Pytorch 1.9.1 (Paszke et al., 2017). The network was optimized based on the evidence lower bound (ELBO) objective by using the reparameterization trick and ADAM optimization (Kingma and Welling, 2013). In our training process, we also injected a unit Gaussian noise *N*(*0,0.01*) to the spectrograms in order to make the network learn robust representations of the motif syllables (Poole et al., 2014). We fixed the latent dimension to 32 for both of our training runs. The approximate posterior was parameterized as a normal distribution with diagonal covariance: *N*(*z*; *μ*, *diag*(*d*)), where *μ* is the latent mean and *d* was the latent diagonal, both of which have a vector of length 32. The VAE architecture was adapted from Goffinet et al, 2021. All activation functions were tanh units. Learning rate was set to 0.001 and batch size was set to 32. The input and output spectrogram size was set to 64 x 256.

### Anomaly Detection Analysis with VAEs

Anomaly detection involves identifying unusual observations that do not conform to the expected behavior (Chalapathy and Chawla, 2019; Chandola et al., 2009; Masaki et al., 2021). To test for the existence of abnormal vocalizations, reconstructed syllable spectrograms from each bird were generated via the VAE trained with sham birds’ syllable spectrograms only. The quality of the reconstruction for each syllable spectrogram was determined via the mean-squared error (MSE):

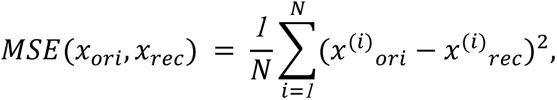

where *x*_*ori*_ is the input syllable spectrogram and *x*_*rec*_ is the reconstructed spectrogram from the VAE decoder, and *N* is the number of pixels. In the juvenile-lesion song analysis, we classify a syllable spectrogram as abnormal if the reconstructed error value exceeds the 99.9th percentile of the sham test set, whereas, in adult-lesion song analysis, we consider a syllable spectrogram to be abnormal if the reconstructed error value exceeds the 99.9th percentile of the pre-lesion test set. In addition, we characterize the MSE distribution tail difference between the *i*th lesion bird and *j*th sham bird in the test set with the following measure:

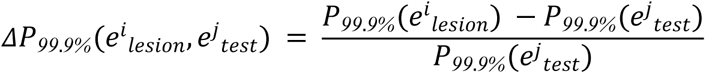

where, *e*^*i*^_*lesion*_ and *e*^*j*^_*test*_ are the mean-squared error distributions for *i*th lesion bird and *j*th sham bird in the test set respectively, and *P*_*99.9%*_(*e*) is the 99.9% percentile of mean-squared error distribution. We then define the anomaly score for the *i*th lesion bird as:

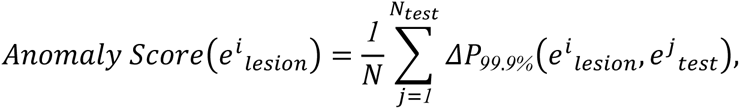

where the *i*th lesion bird is compared to every sham bird in the test set. Similar analysis is performed for the tutor bird and adult birds. In essence, the anomaly score computes how far away in acoustic space the outlying syllables of a bird’s repertoire are from the syllables of the sham-lesioned dataset.

### Principal Component Analysis (PCA)

We complemented our anomaly detection analysis by performing PCA (Jolliffe and Cadima, 2016) on the 32-dimensional latent mean vector obtained via the VAE encoder for each syllable spectrogram. The first three principal components of the juvenile-lesion song capture 97.33% of the variation, and the adult-lesion song’s first three principal components capture 75.37% (for example in Fig 3K and 5F; remaining data shown in Sup Table 1 and 2). The spatial distribution of the abnormal syllables (based on the above cutoff) was then visualized in a two-dimensional projection for the first three components (Figs 3-5, and Sup Figs 3_2 and 5_1).

## Results

### LHb receives inputs from VP and AIV and projects to VTA

Retrograde and anterograde viral tracing experiments were used to characterize LHb connectivity. First, to test for a VP-LHb-VTA pathway, we injected scAAV9-CBh-GFP into VP and retrograde cholera toxin subunit B (CTB) into VTA (n = 9 hemispheres), and we always observed VP axons overlying VTA-projecting cell bodies in LHb (Fig. 1A; n = 9/9 hemispheres). Next, to test for inputs to LHb, and at the same time test if these inputs overlap with inputs to VTA, we injected distinctly colored retrograde tracers into VTA and LHb (n = 6 hemispheres). LHb- and VTA-projecting neurons were observed in VP and the ventral anterior intermediate arcopallium (AIV), part of the auditory pallium known to send projections to STN and VP (Bottjer et al., 2000; Chen et al., 2019; Das and Goldberg, 2022; Mandelblat-Cerf et al., 2014) and also to send singing-related error signals to VTA that control song plasticity (Kearney et al., 2019; Mandelblat-Cerf et al., 2014). We wondered if the populations of neurons in VP and AIV that project to VTA and LHb overlapped or arose from separate populations (as in mammalian BG inputs to LHb) (Knowland et al., 2017; Wallace et al., 2017). Consistent with findings in mammals, we rarely observed overlap in LHb- and VTA-projecting populations in VP and AIV (AIV: Fig. 1B middle, visually estimated at less than 5% overlap in 5/5 hemispheres where cells were observed; VP: Fig. 1B right, LHb neuron labeling seen in 6/6 hemispheres). These results suggest that VP and AIV could send distinct signals to LHb and VTA. Next, to test if LHb axons target the part of the VTA that project to Area X, we injected anterograde virus HSV-mCherry into LHb, and retrograde CTB into Area X. LHb axons overlapped Area X-projecting VTA neurons (Fig. 1C; n = 3/3 hemispheres). Altogether, these data show that LHb receives inputs from two structures important for song learning, VP and AIV, and in turn projects to the part of VTA that sends singing related error signals to Area X. LHb is thus anatomically positioned to contribute to vocal learning.

### VP Microstimulation Activates LHb Neurons

To test for functional connectivity between VP and LHb, specifically VTA projecting LHb neurons (LHb_VTA_), we recorded from LHb neurons in anesthetized birds while electrically stimulating VP (n = 3 birds). A subset of birds (n = 2) were also implanted with stimulation electrodes in VTA to antidromically identify LHb_VTA_ neurons, which were further confirmed with collision testing (n = 2 neurons; Fig. 2A,C). LHb neurons that were not tested for antidromic responses (n = 2), or did not exhibit an antidromic response (n = 4), were classified as LHb_other_ neurons. All LHb neurons recorded were phasically activated by VP stimulation (n = 8/8 neurons; Fig. 2B, Z-tests for significance, Methods). Two LHb_VTA_ neurons exhibited a very short latency response to VP stimulation (0.0025 +/- 0.0035 s; mean +/- SD; Fig. 2D,E; Methods), as well as two LHb_other_ neurons (0.0075 +/- 0.0035 s; tested, not antidromic; not shown). The remaining LHb_other_ neurons showed a slightly longer latency response (0.015 +/- 0.0041 s; n = 4; 2 not tested, 2 tested, not antidromic; Fig. 2F,G; Methods). These results, combined with the anatomical tracing, further support the idea that VP is functionally connected to LHb, and specifically LHb_VTA_ neurons.

### LHb Microstimulation Inhibits VTA DA Neurons

To test if LHb stimulation can influence the activity of Area X projecting VTA neurons (VTA_X_), as well as other local VTA neurons, we recorded VTA neurons in anesthetized birds implanted with bipolar stimulation electrodes in LHb (for orthodromic stimulation) and Area X (for antidromic identification of VTA_X_ neurons) (n = 5 birds; Fig. 2H, Methods). Responses to bursts of LHb stimulation varied across VTA neurons (Fig. 2I), however antidromically identified VTA_X_ neurons consistently exhibited a sustained suppression, or suppression followed by activation (0.010 +/- 0.0089 s; mean +/- SD; n = 6; Fig. 2K-N; Methods), consistent with past work in mammals (Christoph et al., 1986; Ji and Shepard, 2007; Matsumoto and Hikosaka, 2007). VTA neurons not antidromically identified were classified as VTA_other_ neurons; these neurons exhibited low latency activations (0.018 +/- 0.0035 s; n = 2; Fig. 2O-P; Methods), suppressions (0.013 +/- 0.0096 s; n = 6; Fig. 2Q; Methods), or no change (n = 5; not shown) due to LHb burst stimulation. The average spike width, or duration, of VTA_X_ neurons was larger than that of VTA_other_ neurons (Fig. 2J; p = 0.00058, t test; Li et al, 2012). Together, these results show that LHb neurons are able to inhibit VTA_X_ neurons, dopaminergic neurons that encode performance error signals and are vital for song learning, and are functionally connected to VTA_other_ neurons, including putative VTA interneurons that may implement the signal inversion necessary to suppress DA firing.

### LHb Lesions in Juveniles Cause Abnormal Adult Vocalizations

To determine the contributions of LHb signaling to vocal learning and production, we conducted lesion experiments in juvenile birds and recorded their songs in adulthood. All birds were housed with conspecifics and had regular exposure to tutor song until the start of the experiment (37 - 43 dph) when they were acoustically isolated and vocally recorded for the remainder of the experiment. After the onset of babbling, birds were divided into two experimental groups: the lesion group (n = 17 birds) which had bilateral LHb lesions using excitotoxic (2% N-methyl-DL-aspartic acid, NMDA) or electrolytic methods (Methods), and the sham group (n = 9 birds) which had saline injected or electrodes inserted but no current passed (Fig. 3 A-C; Methods). Due to the depth and small size of the LHb, as well as the months between lesion and histology (to study vocal development), we could not confidently quantify lesion size; thus, though lesion protocols were validated in non-experimental birds, the following results should be cautiously interpreted as results from partial LHb lesions (Methods).

We examined the adult songs of birds lesioned as juveniles and found that LHb lesions did not have a significant effect on the capacity of birds to copy tutor song syllables (tutor imitation score, p = 0.71, unpaired, two-sided Wilcoxon rank sum test, n = 17 lesion and n = 9 sham; Methods). A subset of LHb lesion birds produced a good copy of their tutor song, comparable or better than shams (Fig. 3G). However, roughly half of the lesioned birds produced highly abnormal syllables in addition to their normal-appearing ones (Fig. 3D-F). As evident by the example spectrograms presented in Fig. 3D-F (bottom three rows) and Fig. 4A-C (top two rows), many LHb lesion birds produced species atypical vocalizations within song bouts, as part of their motif, and as isolated calls. Additionally, for the same dataset, lesion birds produced vocalizations that were roughly 400 ms longer than vocalizations seen in either the sham or tutor birds (99.9th percentile syllable duration for tutor: 280 ms, for sham: 313 ms, and for lesion: 734 ms; sham - tutor p = 3.48e-18; sham-lesion p = 5.35e-197 and tutor-lesion p = 5.81e-7, pairwise Kolmogorov-Smirnov tests with Bonferroni correction; Sup Fig 3_1F). These abnormally long vocalizations also often contained high-pitched, narrow frequency bands that repeated, strangely resembling canary trills or starling buzzes (examples in Fig. 3E and Fig. 4A) (Henry et al., 2015; Williams, 2004). In other birds, abnormalities included: shorter-duration unusual ‘elements’ added to ‘typical’ syllables, and rare ‘Wsst’-type syllables described as a ‘hissing sound’, similar to vocalizations often used before one finch attacks another (examples in Fig. 3D, Fig. 4B,C, and Sup Fig 3_1) (Zann, 1996). These abnormal vocalizations tended not to crystalize (e.g. the bird shown in Fig. 4C produced abnormal syllables with variable durations; Sup Fig 3_1).

Many birdsong segmentation algorithms have been developed to cluster and quantify differences between cleanly separable song syllables (Cohen et al., 2022; Goffinet et al., 2021; Sainburg et al., 2020). Here we aimed to quantify the production of rarely produced, atypical syllables that lacked a stereotyped acoustic structure within and across birds, i.e. to systematically detect anomalous vocalizations. To address this problem, we used anomaly detection methods previously used in biomedical and surveillance applications but not, to our knowledge, for the detection of strange vocal events (Chalapathy and Chawla, 2019; Chandola et al., 2009; Masaki et al., 2021). To do so, we leveraged Variational AutoEncoders (VAEs) (Goffinet et al., 2021; Kingma and Welling, 2013; Sainburg et al., 2020) to learn a reduced-dimensional and informative representation of ‘normal’ vocalizations from a subset of adult sham-lesioned birds’ undirected songs (n = 5 birds; Methods). In order to acquire the data to train the neural network, we first hand-segmented syllables across five days for each lesion and sham bird, and one day for tutor birds (90 - 94 dph lesion and sham; n = 17 lesion; n = 9 sham; n = 12 tutors; Methods). We then performed UMAP (uniform manifold approximation and projection) on each bird to obtain a low-dimensional visualization for every vocalization (analyzed in the form of a spectrogram). The spectrograms were clustered in an unsupervised manner yielding 4-7 clusters from each bird (Sup. Fig. 3_2; Methods), roughly forming one cluster for each song syllable, call type, or abnormal vocalization (Sainburg et al., 2020). Next, we selected five of the nine sham birds to train a VAE to learn a low-dimensional representation of species typical vocalizations from control birds. We repeated this procedure with six randomly selected subsets of the sham birds to prevent selection bias and to produce distinct networks trained on varying selections of normal bird songs (Methods).

Each trained network provided two outputs for each input spectrogram: 1) a low dimensional representation (32 dimensional vector), and 2) a faithful reconstruction with low mean-squared error when compared to the original ‘normal’ syllables, i.e with low reconstruction error values (Fig. 3H, black and gray; Methods). We next compared the reconstruction errors between the LHb lesion vocalizations to the four held-out sham birds. If anomalous syllables were equally distributed between sham and lesion birds, then high reconstruction errors would be observed in both groups. Alternatively, if lesioned birds were significantly more likely to contain atypical syllables, then their reconstruction error distributions would exhibit long tails at high values. We observed that the LHb lesion bird data indeed exhibited significantly larger tails in the error distributions (Fig 3H, red). These abnormal syllables in all datasets were identified by using the 99.9th percentile of data in the sham bird test set (Fig. 3H, gray) as the threshold: all reconstruction error values higher than this threshold were considered abnormal vocalizations (Fig. 3H; red vertical line). The vast majority of abnormal vocalizations were produced by the LHb lesion birds (red: abnormal, consisting of 9.06% of total lesion bird vocalizations; blue: normal; Fig. 3H,I; Sup Table 1), and very few, if any, were found in the other datasets (0.125% sham abnormal vocalizations; 0.52% tutor abnormal vocalizations; Sup Table 1). We further characterized the error distribution’s tail at the bird level across the six replicate VAEs and found the LHb lesion birds were significantly more likely to contribute to outlying syllables than the sham and tutor birds (Averaged Anomaly Score; Kruskal-Wallis test with Bonferroni method multiple comparison, α = 0.05; n = 12 tutor, n = 9 sham, and n = 17 lesion birds; Fig 3J).

We then made use of the low dimensional representation of each vocalization from the VAE to test if representations for abnormal syllables were different from normal syllables. We performed Principal Component Analysis (PCA) on the 32-dimensional output vector of the VAE (Methods). We used the first three principal components to visualize the vocalizations. The LHb lesion birds’ abnormal syllables occupied a distinct region in PC space as compared to the tutors’ and sham pupils’ motif syllables (1st row: tutor; 2nd row: sham; and 3rd row: lesion; Fig 3K; two additional permutations shown in Sup Fig 3_3). Together, these analyses show that LHb lesions in juvenile finches can cause vocal abnormalities in adult undirected song.

### LHb lesion-associated vocal abnormalities are produced in female-directed courtship song

Male zebra finches will spontaneously sing while alone, with the goal of learning or maintaining a song similar to that of their ‘template’ or tutor song memory (Zann, 1996). When presented with a female, males transition to a courtship state and female-directed song (Kao et al., 2008). We presented LHb lesion and sham birds (n = 17 lesion, n = 4 sham) with a female to test if courtship singing also contained abnormal acoustic elements (91-107 dph; Fig. 4). All males readily and robustly sang female-directed song, and lesion birds whose undirected song contained abnormal vocalizations exhibited the same types of species atypical vocalizations in female-directed song (Fig. 4A-C and Sup Fig 4_1 second column). Two birds in particular produced a more exaggerated version of their trilled note in female-directed song (Fig. 4A-B, also shown in Sup Fig 4_1B,D).

We set out to compare the reduced representations and reconstructed spectrograms for female-directed song between LHb lesion and sham birds using the same methods and trained network used for undirected song (Methods; Fig. 4). Again, we refer to vocalizations with reconstruction error greater than the 99.9th percentile of the sham test set as abnormal syllables (Fig. 4E, red vertical line; refer to Fig. 4F for example spectrograms and reconstructed spectrograms from the VAE). We again observed that the LHb lesion bird spectrograms had a significantly larger tail in the error distribution (4.68% of lesion bird vocalizations were abnormal) as compared to sham bird female-directed song (0.65% of sham vocalizations were abnormal; Fig. 4E and Sup Table 1). At the individual bird level, we found again the LHb lesion birds’ female directed song has a significantly different distribution tail as compared to sham birds’ (Averaged Anomaly Score; Kruskal-Wallis test with Bonferroni method multiple comparison: α = 0.05; n = 9 sham undirected, n = 17 lesion directed and n = 9 sham directed).

To further visualize the differences in vocalizations, we projected the reduced representation of female-directed song syllables from the VAE onto the PC space created with undirected song (Methods). This showed that the female-directed song exhibited abnormal vocalizations that occupied a similar region as the abnormal vocalizations during undirected song (Fig. 3K, bottom row; Fig 4G, bottom row); these outlying clusters were not observed in the sham bird data (Fig. 4G, top row). Thus, LHb lesioned birds sang abnormal songs even to females.

### LHb Lesions in Adults Do Not Alter Vocalizations

In songbirds, lesioning the basal ganglia in juveniles impairs song learning but lesions in adults do not significantly impair production of the learned song (Bottjer et al., 1984; Scharff and Nottebohm, 1991). To test if LHb lesions impair adult song production, we lesioned LHb in crystalized adults (>150 dph; Methods). To compare vocalizations before and after bilateral LHb lesion, adult lone song was recorded for at least 5 days prior to lesion, and at least 60 days post-lesion, as well as one session of female-directed song (n = 5 adult birds; 57-84 days post-lesion). As evident by the example spectrograms from three lesioned adults, vocalizations were not altered by LHb lesions, even up to two months post-lesion (Fig. 5 A-C). To compare motif similarity pre and post-lesion in lone song, we computed the similarity score between: 1) two days pre-lesion (2-3 days apart; Fig. 5D, black center), 2) pre-lesion and 60 days post-lesion (Fig. 5D, red), and 3) control adult bird song recorded 60 days apart (Fig. 5D, black left) and found no significant difference between any of the three conditions (Glaze and Troyer, 2013; Hyland Bruno and Tchernichovski, 2019).

To further verify the absence of abnormal syllables in undirected and female-directed post-lesion songs, we trained a new network based on the pre-lesion lone song vocalizations (n = 3 adult birds). Using the same method as the juvenile LHb lesion birds, we computed the reconstruction error values for the pre-lesion undirected song test set (n = 2 birds), post-lesion undirected, and female-directed song (n = 5 birds; Methods). We found that only 1.23% undirected and 2.94% of female-directed song vocalizations were above the 99.9th percentile threshold (Fig 5E; Sup Table 2). We found no significant difference in the distribution tails for adults (Averaged Anomaly Score; Kruskal-Wallis test p = 0.9139; n = 5 adult pre-lesion, n = 5 adult post-lesion directed and n = 5 adult post-lesion directed; Fig 5F). For completeness, we also performed PCA analysis and did not observe any significant differences between the reduced representation of pre-lesion, post-lesion, and female-directed vocalizations (Fig 5G and Sup Fig 5_1). Therefore, the LHb lesions in adults do not impair crystalized song production or maintenance over month timescales.

## Discussion

**In a tennis game,** double faulting or acing an opponent on match point causes deep sadness or elation. In 1986, Vernon Brooks proposed the ‘limbic comparator hypothesis’ in which neural circuits underlying the full range of emotional qualia are used in the service of sensorimotor learning (Brooks, 1986). Here we tested this idea in songbirds by focusing on the lateral habenula, an evolutionarily conserved input to midbrain dopamine neurons implicated in reward processing, reinforcement learning, depression and learned helplessness - but not, to our knowledge, in an intrinsically motivated motor learning task like birdsong (Andalman et al., 2019; Boulos et al., 2017; Namboodiri et al., 2016; Ullsperger and von Cramon, 2003). The recent discovery of dopaminergic mechanisms for birdsong learning led us to hypothesize that the LHb may be part of a song evaluation system that includes regions of the songbird brain outside the classic, nucleated song system (Chen et al., 2019; Chen and Goldberg, 2020; Chung and Bottjer, 2022; Das and Goldberg, 2022; Gale and Perkel, 2010; Hisey et al., 2018; Mandelblat-Cerf et al., 2014). Through the combination of anatomical and functional mapping techniques, we discovered that LHb receives inputs from VP and AIV, two brain regions important for song learning. We also discovered that LHb projects to the part of VTA that projects to Area X, and that LHb stimulation suppresses the activity of VTAx DA neurons. Finally, we discovered LHb lesions in juvenile birds results in adults that can still imitate tutors but exhibit strange, high-pitch and long duration atypical vocalizations that were produced during both undirected and courtship song.

The significance of the strange vocalizations produced by LHb lesioned birds is unclear, but we note that variable, unstructured and long-duration high pitch notes have also been observed in socially isolated zebra finches raised without male tutors (Feher et al., 2009; Williams et al., 1993) and birds that had DA antagonist infused into HVC during tutoring (Tanaka et al., 2018). Yet unlike isolate birds, LHb lesion birds that produced these strange syllables were still able to imitate tutor song and produce structured song motifs with regular rhythms (Fig 3-4 and Sup Fig 4_1). However, they would place these isolate-like vocalizations prior to, within, or after their stereotyped motif, causing irregularities in bout structure in both undirected and female-directed song (Sup Fig 4_1). Also, LHb lesion birds sometimes uttered the abnormal vocalizations more like a call – as single vocalizations surrounded by hundreds of milliseconds to seconds of silence (Fig 3 and 4, Sup Fig 3_1 and 4_1). Further, no alterations to motif or vocalizations were found in adult LHb lesion birds, supporting the idea that LHb plays a role in learning vocalizations, but not maintenance.

An important caveat of our study is that, though we carefully calibrated our electrolytic and chemical lesion methods, and always confirmed substantial (>50% of tissue by area) habenular tissue damage in lesioned birds, because of the months between lesion and histology we were not always able to obtain high quality, cellular resolution estimates of lesion size and scope. Thus we cannot rule out that our lesions also affected medial habenula, a cholinergic nucleus that has projections to thalamic areas that, in turn, may project to HVC (Akutagawa and Konishi, 2005). Future studies recording from, or optogenetically manipulating, VTA projecting LHb neurons in freely singing birds with auditory feedback would definitively test how LHb plays a role in song evaluation, and what error- or song-related signals LHb sends to dopaminergic midbrain neurons.

---

**Supplementary Figure 3_1.**
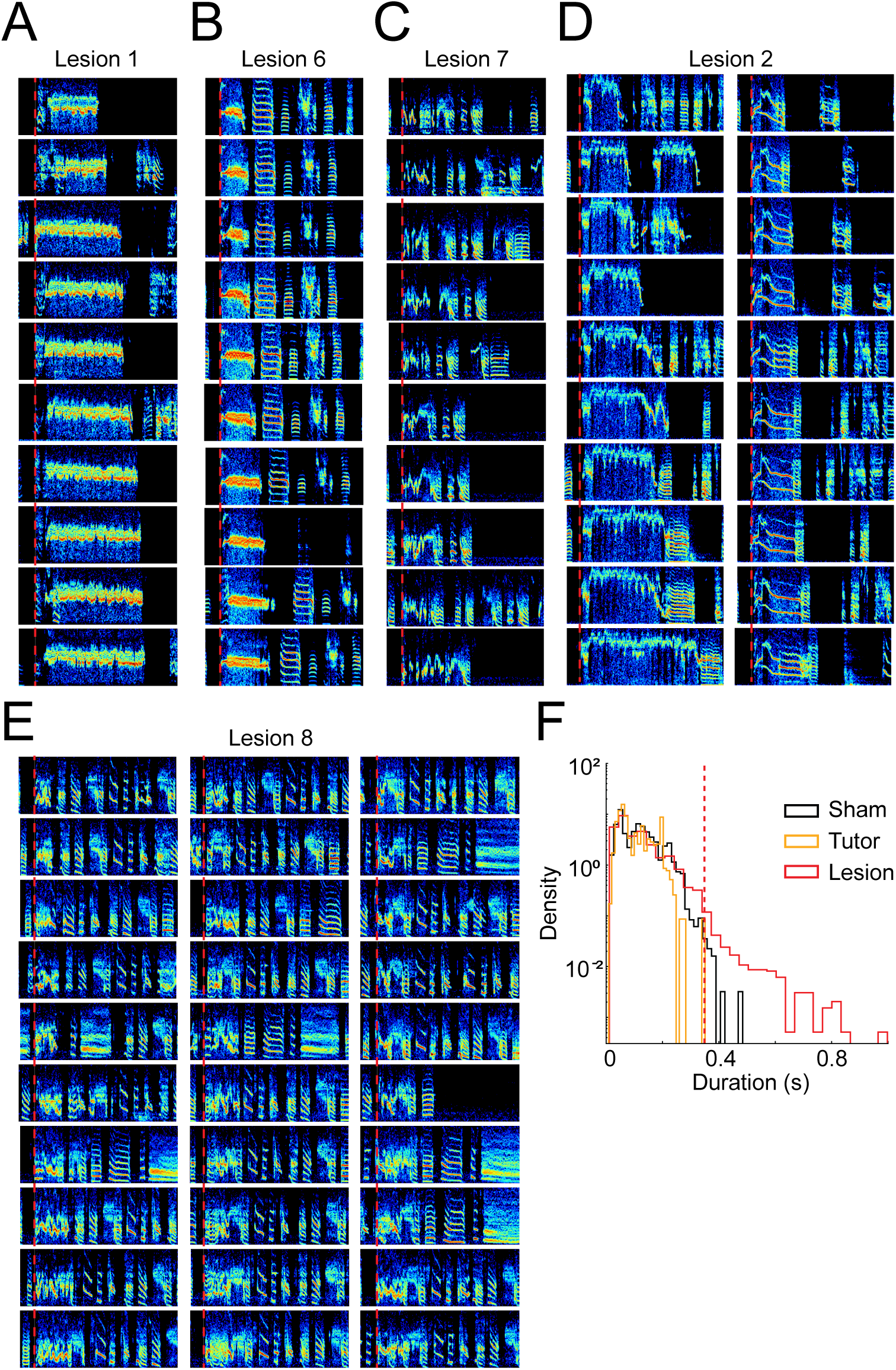
(with Figure 3). Abnormal vocalizations can exhibit long and variable duations, evident as long tails in the syllable duration distribution. A: example undirected song spectrograms from adult birds that experienced an LHb lesion as a juvenile. Vocalizations were recorded at 92 dph and illustrate the distribution of durations for this single syllable type (bird also shown in Fig. 3D). All spectrograms are 1 second in duration; red dashed line denotes abnormal syllable onset. B: same as in A, except for the LHb lesioned bird shown in Fig. 4C. C: same as A for a bird not previously shown. D: same as A, except each column depicts a different vocalization from bird shown in Fig. 3E. E: same as D, except bird not previously shown. F: the distribution of sham, tutor, and lesion bird syllable durations during adult undirected song (black: sham; yellow: tutor; lesion: red). Pairwise Kolmogorov-Smirnov tests with Bonferroni correction were performed for statistical differences between the distributions (sham - tutor p = 3.48e-18; sham-lesion p = 5.35e-197 and tutor-lesion p = 5.81e-7).

**Supplementary Figure 3_2.**
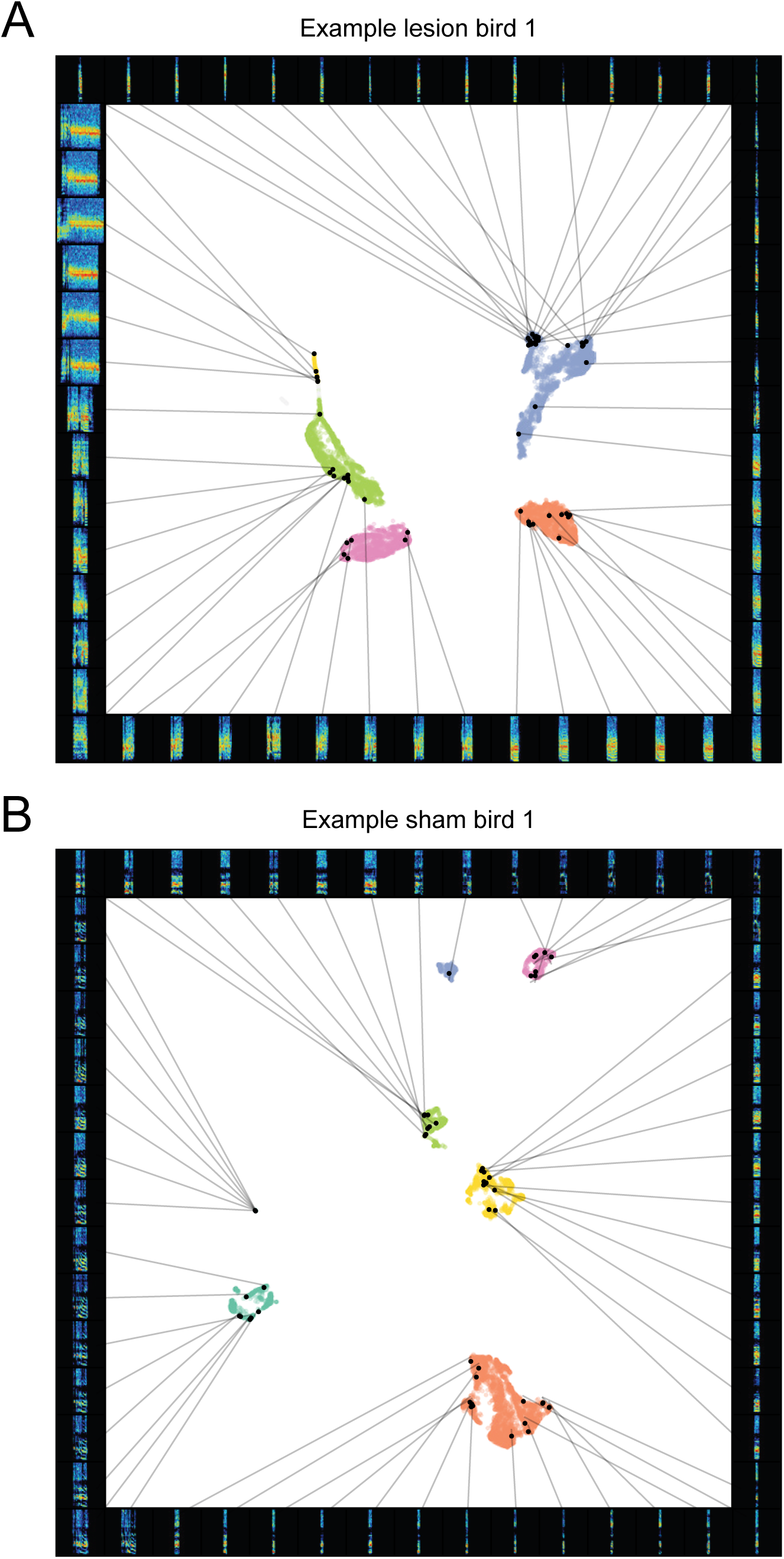
(with Figure 3). Uniform Manifold Approximation and Projection (UMAP) for example LHb lesion and sham birds. A: example UMAP projection for the example LHb lesion bird shown in Fig 3D. Each cluster is differently colored, and corresponds to a distinct syllable. Example spectrograms from each cluster are populated around the UMAP projection. B: data same as A but for example sham bird seen in Fig 3G.

**Supplementary Figure 3_3.**
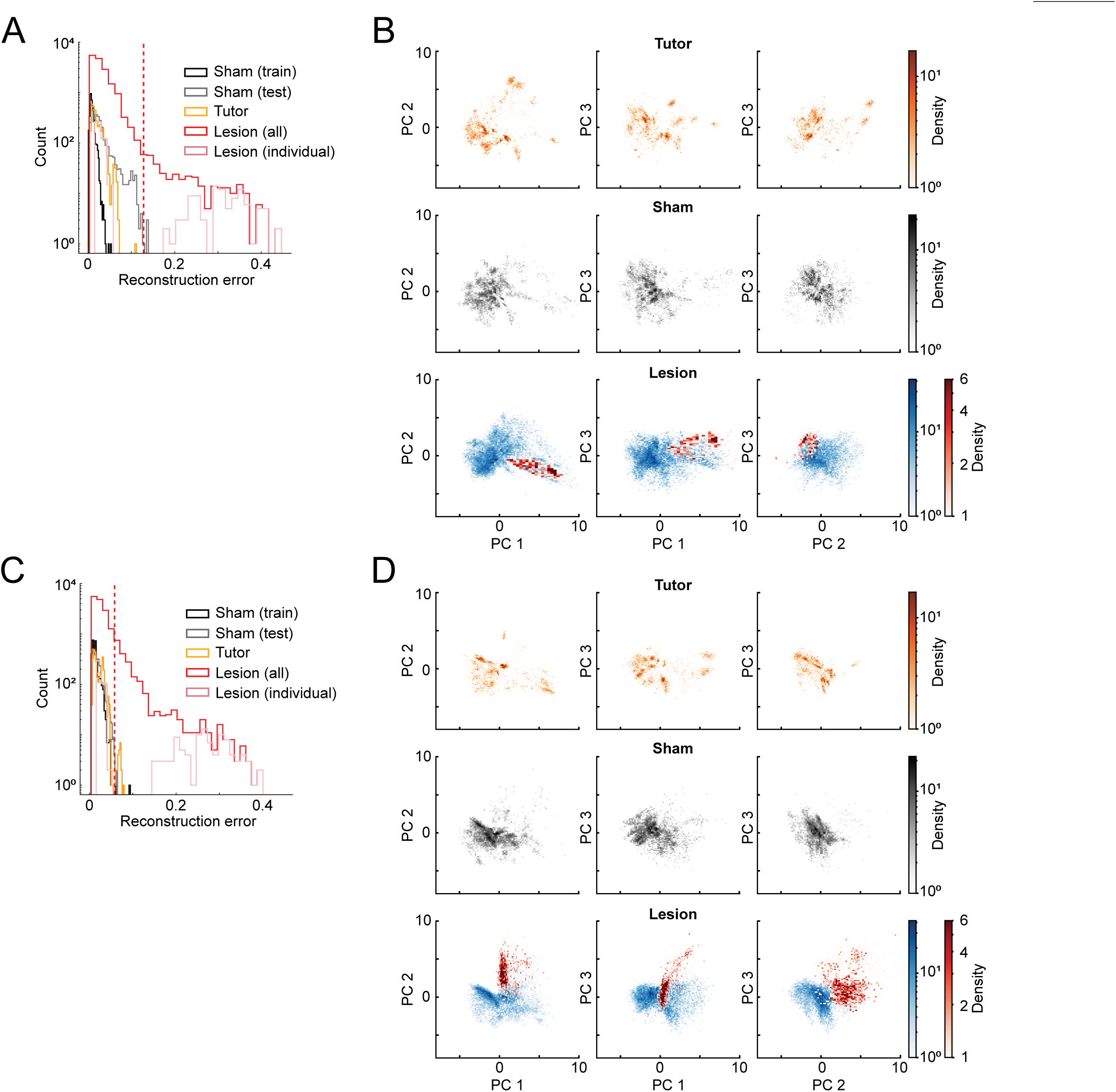
(with Figure 3). Identification of abnormal syllables in LHb lesioned birds is robust across independent replicate VAEs trained with different subsets of sham birds. A: reconstruction error distribution for all datasets. The red vertical line corresponds to the threshold set at the 99.9th percentile of sham bird (test set) vocalizations, which is used to divide the lesion dataset into normal and abnormal vocalizations. Sham birds (training set: black outline; test set: gray), LHb lesion birds (average: red; individual bird shown in 3D: pink), and tutor bird (yellow) vocalizations. B: two-dimensional projection of the PC space for the first three components. Top row corresponds to tutor vocalizations, middle row corresponds to all sham bird vocalizations (training and test set), and bottom row corresponds to LHb lesioned birds vocalizations (blue: normal syllables; red: abnormal syllables as determined by the threshold in A). C-D: same as A-B with an additional cross validation example from a VAE independently trained on a distinct subset of sham birds.

**Supplementary Figure 4_1.**
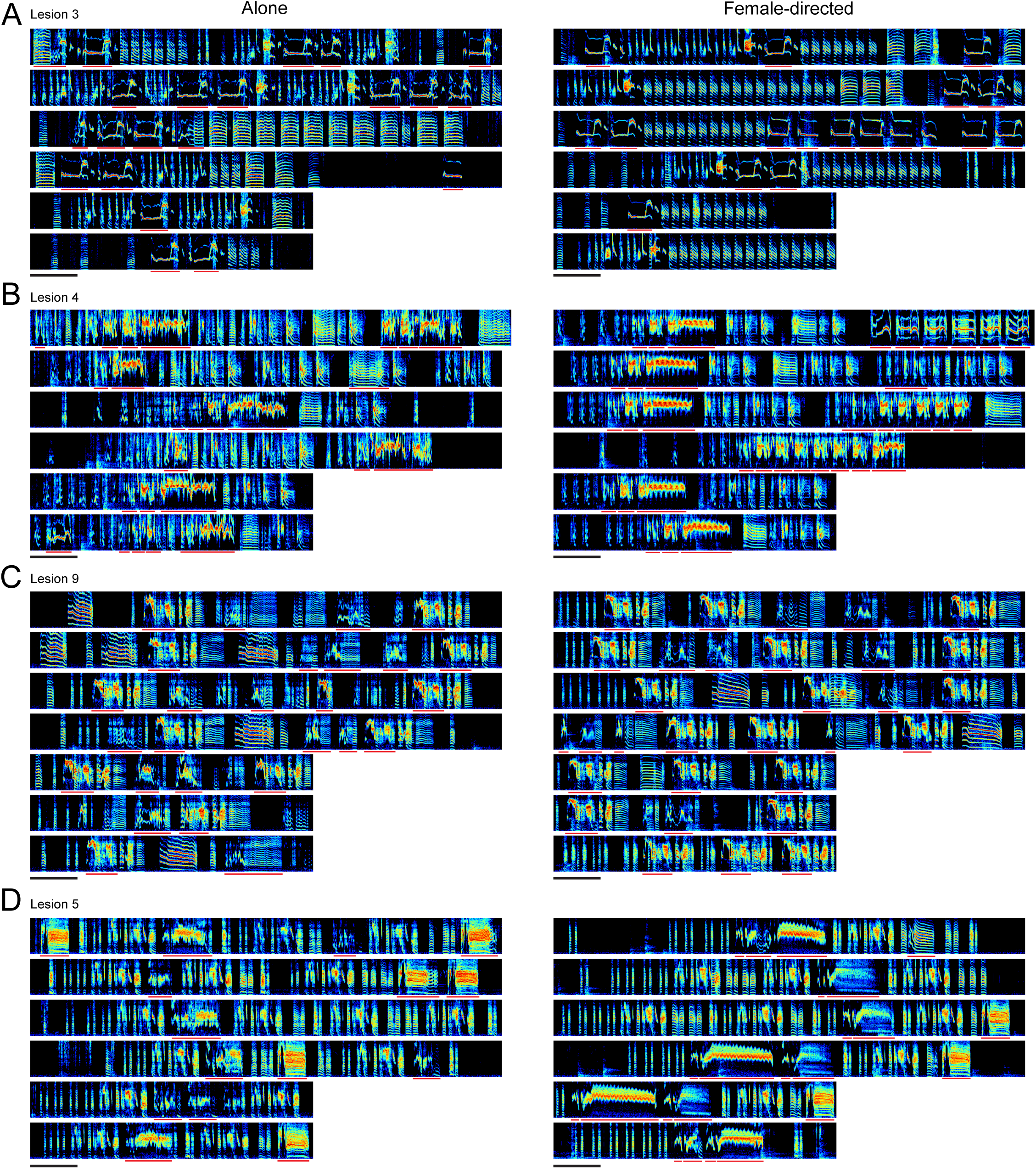
(with Figure 4). Abnormal vocalizations are produced in both undirected and female-directed courtship songs. A: example song and calls from a single bird (left: undirected song; right: female-directed song; same bird as Fig. 3F). Scale bar is 500 ms. Red underlines indicate abnormal vocalizations. B: same as in A, except the bird shown in Fig. 4A. C: same as in A, except the bird was not previously shown. D: same as in A, except the bird previously shown in Fig. 4B.

**Supplementary Figure 5_1.**
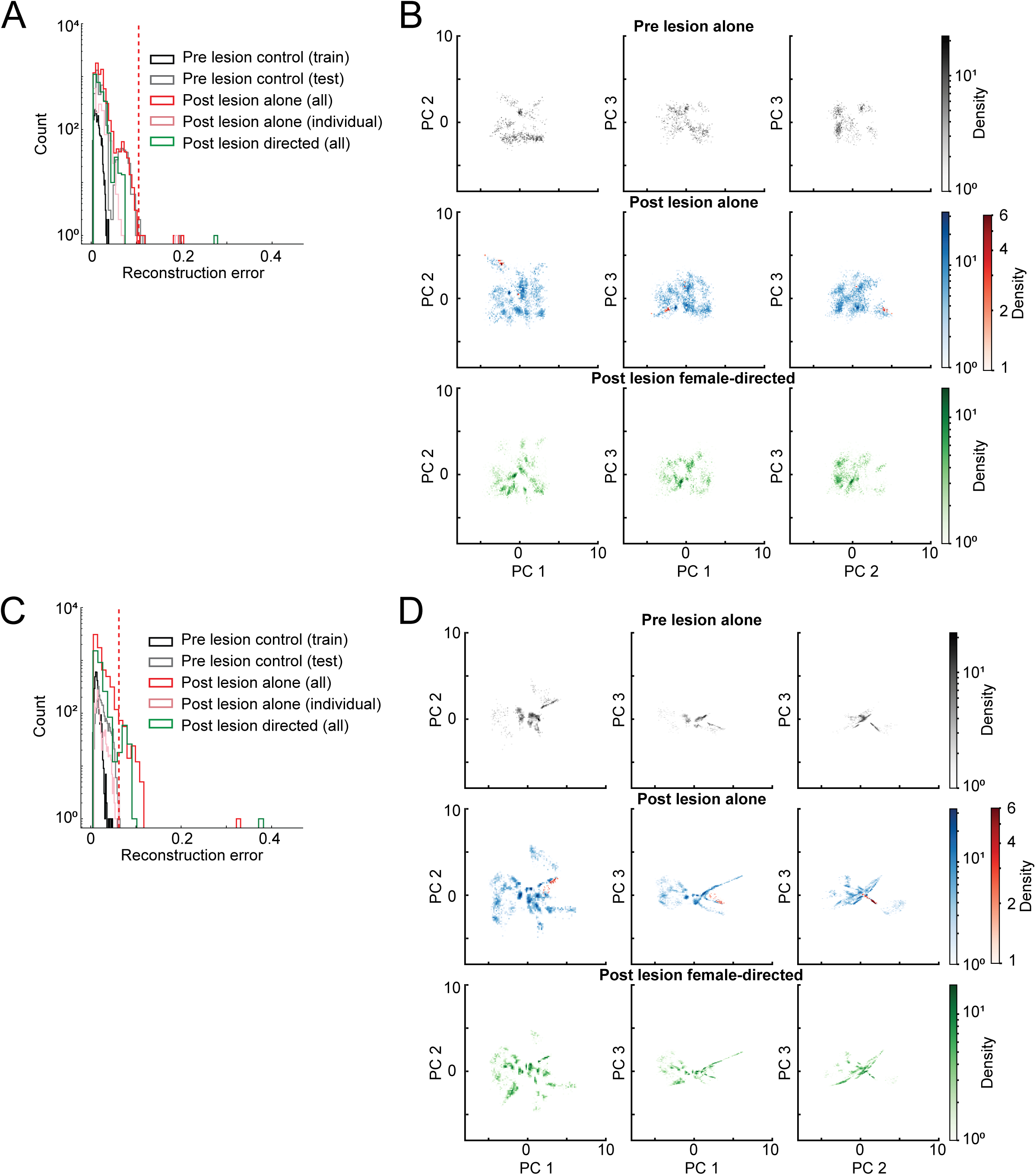
(with Figure 5). Results from figure 5 are robust across replicate VAEs trained with different subsets of sham birds. A: reconstruction error distribution for all datasets. The red vertical line corresponds to the threshold set at the 99.9th percentile of the test set (pre-lesion control) vocalizations, which is used to divide the post-lesion dataset into normal and abnormal vocalizations. Pre-lesion data (training set: black outline; test set: gray), post LHb lesion undirected data (average: red; individual bird shown in Fig. 5A: pink), and post LHb lesion female-directed data (green) vocalizations. B: two-dimensional projection of the PC space for the first three components. Top row corresponds to pre-lesion undirected vocalizations, middle row corresponds to post-lesion undirected (blue: normal syllables; red: abnormal syllables as determined by the threshold in A), and bottom row corresponds to post-lesion female directed vocalizations. C-D: same as A-B except another cross validation analysis example.

**Supplementary Table 1.** (with Figs 3 and 4). Variational Autoencoder (VAE) results for six different runs where the sham juvenile birds were permuted between training and test sets.

| Training Set | Training Set Size | Undirected Reconstruction Error<br>>99.9th pct sham test |  | Directed Reconstruction Error<br>>99.9th pct sham test | PCA Explained Variance<br>(First Three Components) |
| --- | --- | --- | --- | --- | --- |
| 1 | 4700 | Tutor Lesion | 0.52%<br>9.06% | Directed Sham 0.65%<br>Directed Lesion 4.68% | 97.33% |
| 2 | 5000 | Tutor Lesion | 0%<br>1.99% | Directed Sham 0%<br>Directed Lesion 0.7% | 96.6% |
| 3 | 3300 | Tutor Lesion | 0.74%<br>11.71% | Directed Sham 0.81%<br>Directed Lesion 5.63% | 71.34% |
| 4 | 3200 | Tutor Lesion | 0.71%<br>13.25% | Directed Sham 1.05%<br>Directed Lesion 6.11% | 73.64% |
| 5 | 3600 | Tutor Lesion | 0.0%<br>7.90% | Directed Sham 0.48%<br>Directed Lesion 3.14% | 66.35% |
| 6 | 5100 | Tutor Lesion | 0.0%<br>6.64% | Directed Sham 0.57%<br>Directed Lesion 3.16% | 80.34% |

**Supplementary Table 2.** (with Fig. 5). Variational Autoencoder (VAE) results for six different runs where the adult birds were permuted between training and test sets.

| Training Set | Training Set Size | Reconstruction Error<br>>99.9th pct pre-lesion test |  | PCA Explained Variance<br>(First Three Components) |
| --- | --- | --- | --- | --- |
| 1 | 4000 | Directed Lesion | 3.49%<br>2.36% | 75.37% |
| 2 | 3800 | Directed Lesion | 1.11%<br>0.83% | 61.37% |
| 3 | 3400 | Directed Lesion | 3.53%<br>2.71% | 96.04% |
| 4 | 3600 | Directed Lesion | 2.28%<br>2.79% | 71.33% |
| 5 | 4000 | Directed Lesion | 4.45%<br>1.77% | 90.03% |
| 6 | 4200 | Directed Lesion | 2.42%<br>0.21% | 90.03% |

## Acknowledgments

We thank Archana Podury and Kamal Maher for help with anatomical tract tracing. We thank Nao Uchida and members of the Goldberg lab for comments on the project and the manuscript.

## Grants

This work was supported by the NIH R01NS094667 (JHG), NIH R01NS116595 (HKT and IC) and a Mong Cornell Neurotech seed grant (AR and HKT).

## Disclosures

None

## Author Contributions

AR and JG designed the experiments. AR and RC performed the experiments. AR, JG and HKT analyzed the data. HKT was advised by IC. AR, JG and HKT wrote the manuscript.

## Notes

### Competing Interest Statement

The authors have declared no competing interest.

## References

Akutagawa E, Konishi M. 2005. Connections of thalamic modulatory centers to the vocal control system of the zebra finch. Proc Natl Acad Sci U S A 102:14086–14091.

Andalman AS, Burns VM, Lovett-Barron M, Broxton M, Poole B, Yang SJ, Grosenick L, Lerner TN, Chen R, Benster T, Mourrain P, Levoy M, Rajan K, Deisseroth K. 2019. Neuronal Dynamics Regulating Brain and Behavioral State Transitions. Cell 177:970–985 e20.

Bottjer SW, Brady JD, Cribbs B. 2000. Connections of a motor cortical region in zebra finches: relation to pathways for vocal learning. J Comp Neurol 420:244–260.

Bottjer SW, Miesner EA, Arnold AP. 1984. Forebrain lesions disrupt development but not maintenance of song in passerine birds. Science 224:901–903.

Boulos L-J, Darcq E, Kieffer BL. 2017. Translating the Habenula—From Rodents to Humans. Biol Psychiatry 81:296–305.

Brooks VB. 1986. How does the limbic system assist motor learning? A limbic comparator hypothesis. Brain Behav Evol 29:29–53.

Chalapathy R, Chawla S. 2019. Deep Learning for Anomaly Detection: A Survey. *arXiv [csLG]*. Chandola V, Banerjee A, Kumar V. 2009. Anomaly detection: A survey. ACM Comput Surv 41:1–58.

Chen R, Goldberg JH. 2020. Actor-critic reinforcement learning in the songbird. Curr Opin Neurobiol 65:1–9.

Chen R, Puzerey PA, Roeser AC, Riccelli TE, Podury A, Maher K, Farhang AR, Goldberg JH. 2019. Songbird Ventral Pallidum Sends Diverse Performance Error Signals to Dopaminergic Midbrain. Neuron 103:266–276 e4.

Christoph GR, Leonzio RJ, Wilcox KS. 1986. Stimulation of the lateral habenula inhibits dopamine-containing neurons in the substantia nigra and ventral tegmental area of the rat. J Neurosci 6:613–619.

Chung JH, Bottjer SW. 2022. Developmentally regulated pathways for motor skill learning in songbirds. J Comp Neurol 530:1288–1301.

Cohen Y, Nicholson DA, Sanchioni A, Mallaber EK, Skidanova V, Gardner TJ. 2022. Automated annotation of birdsong with a neural network that segments spectrograms. Elife 11. doi:10.7554/eLife.63853

Costa RM. 2007. Plastic corticostriatal circuits for action learning: what’s dopamine got to do with it? Ann N Y Acad Sci 1104:172–191.

Das A, Goldberg JH. 2022. Songbird subthalamic neurons project to dopaminergic midbrain and exhibit singing-related activity. J Neurophysiol 127:373–383.

Ding L, Perkel DJ. 2004. Long-term potentiation in an avian basal ganglia nucleus essential for vocal learning. J Neurosci 24:488–494.

Duffy A, Latimer KW, Goldberg JH, Fairhall AL, Gadagkar V. 2022. Dopamine neurons evaluate natural fluctuations in performance quality. Cell Rep 38:110574.

Elston TW, Bilkey DK. 2017. Anterior Cingulate Cortex Modulation of the Ventral Tegmental Area in an Effort Task. Cell Rep 19:2220–2230.

Feher O, Wang H, Saar S, Mitra PP, Tchernichovski O. 2009. De novo establishment of wild-type song culture in the zebra finch. Nature 459:564–568.

Gadagkar V, Puzerey PA, Chen R, Baird-Daniel E, Farhang AR, Goldberg JH. 2016. Dopamine neurons encode performance error in singing birds. Science 354:1278–1282.

Gale SD, Perkel DJ. 2010. A basal ganglia pathway drives selective auditory responses in songbird dopaminergic neurons via disinhibition. J Neurosci 30:1027–1037.

Glaze CM, Troyer TW. 2013. Development of temporal structure in zebra finch song. J Neurophysiol 109:1025–1035.

Goffinet J, Brudner S, Mooney R, Pearson J. 2021. Low-dimensional learned feature spaces quantify individual and group differences in vocal repertoires. Elife 10. doi:10.7554/eLife.67855

Henry L, Craig AJFK, Lemasson A, Hausberger M. 2015. Social coordination in animal vocal interactions. Is there any evidence of turn-taking? The starling as an animal model. Front Psychol 6:1416.

Hisey E, Kearney MG, Mooney R. 2018. A common neural circuit mechanism for internally guided and externally reinforced forms of motor learning. Nat Neurosci 1.

Hoffmann LA, Saravanan V, Wood AN, He L, Sober SJ. 2016. Dopaminergic Contributions to Vocal Learning. J Neurosci 36:2176–2189.

Hyland Bruno J, Tchernichovski O. 2019. Regularities in zebra finch song beyond the repeated motif. Behav Proc 163: 53--59.

Ji H, Shepard PD. 2007. Lateral Habenula Stimulation Inhibits Rat Midbrain Dopamine Neurons through a GABAA Receptor-Mediated Mechanism. J Neurosci 27:6923–6930.

Kao MH, Wright BD, Doupe AJ. 2008. Neurons in a forebrain nucleus required for vocal plasticity rapidly switch between precise firing and variable bursting depending on social context. J Neurosci 28:13232–13247.

Kearney MG, Warren TL, Hisey E, Qi J, Mooney R. 2019. Discrete Evaluative and Premotor Circuits Enable Vocal Learning in Songbirds. Neuron. doi:10.1016/j.neuron.2019.07.025

Kingma DP, Welling M. 2013. Auto-Encoding Variational Bayes. arXiv [statML].

Knowland D, Lilascharoen V, Pacia CP, Shin S, Wang EH-J, Lim BK. 2017. Distinct Ventral Pallidal Neural Populations Mediate Separate Symptoms of Depression. Cell 170:284– 297.e18.

Mandelblat-Cerf Y, Las L, Denisenko N, Fee MS. 2014. A role for descending auditory cortical projections in songbird vocal learning. Elife 3. doi:10.7554/eLife.02152

Marler P. 1997. Three models of song learning: evidence from behavior. J Neurobiol 33:501– 516.

Marler P. 1970. A comparative approach to vocal learning: Song development in white-crowned sparrows. J Comp Physiol Psychol 71:1–25.

Masaki A, Nagumo K, Lamsal B, Oiwa K, Nozawa A. 2021. Anomaly detection in facial skin temperature using variational autoencoder. Artif Life Robot 26:122–128.

Matsumoto M, Hikosaka O. 2009. Representation of negative motivational value in the primate lateral habenula. Nat Neurosci 12:77–84.

Matsumoto M, Hikosaka O. 2007. Lateral habenula as a source of negative reward signals in dopamine neurons. Nature 447:1111–1115.

McInnes L, Healy J, Astels S. 2017. hdbscan: Hierarchical density based clustering. J Open Source Softw 2:205.

McInnes L, Healy J, Melville J. 2018. UMAP: Uniform Manifold Approximation and Projection for Dimension Reduction. arXiv [statML].

Namboodiri VMK, Rodriguez-Romaguera J, Stuber GD. 2016. The habenula. Curr Biol 26:R873–R877.

Paszke A, Gross S, Chintala S, Chanan G, Yang E, DeVito Z, Lin Z, Desmaison A, Antiga L, Lerer A. 2017. Automatic differentiation in PyTorch.

Person AL, Gale SD, Farries MA, Perkel DJ. 2008. Organization of the songbird basal ganglia, including area X. J Comp Neurol 508:840–866.

Poole B, Sohl-Dickstein J, Ganguli S. 2014. Analyzing noise in autoencoders and deep networks. arXiv [csNE].

Reiner A. 2009. You cannot have a vertebrate brain without a basal gangliaThe Basal Ganglia IX, 1. Springer New York. pp. 3–24.

Reiner A, Laverghetta AV, Meade CA, Cuthbertson SL, Bottjer SW. 2004. An immunohistochemical and pathway tracing study of the striatopallidal organization of area X in the male zebra finch. J Comp Neurol 469:239–261.

Sainburg T, Thielk M, Gentner TQ. 2020. Finding, visualizing, and quantifying latent structure across diverse animal vocal repertoires. PLoS Comput Biol 16:e1008228.

Saravanan V, Hoffmann LA, Jacob AL, Berman GJ, Sober SJ. 2019. Dopamine depletion affects vocal acoustics and disrupts sensorimotor adaptation in songbirds. eNeuro 6.

Scharff C, Nottebohm F. 1991. A comparative study of the behavioral deficits following lesions of various parts of the zebra finch song system: implications for vocal learning. J Neurosci 11:2896–2913.

Schultz W, Dayan P, Montague PR. 1997. A neural substrate of prediction and reward. Science 275:1593–1599.

Swanson LW. 2000. Cerebral hemisphere regulation of motivated behavior. Brain Res 886:113– 164.

Tanaka M, Sun F, Li Y, Mooney R. 2018. A mesocortical dopamine circuit enables the cultural transmission of vocal behaviour. Nature 563:117–120.

Tchernichovski O, Nottebohm F, Ho CE, Pesaran B, Mitra PP. 2000. A procedure for an automated measurement of song similarity. Anim Behav 59:1167–1176.

Tian J, Uchida N. 2015. Habenula Lesions Reveal that Multiple Mechanisms Underlie Dopamine Prediction Errors. Neuron 87:1304–1316.

Ullsperger M, von Cramon DY. 2003. Error Monitoring Using External Feedback: Specific Roles of the Habenular Complex, the Reward System, and the Cingulate Motor Area Revealed by Functional Magnetic Resonance Imaging. J Neurosci 23:4308–4314.

Wallace ML, Saunders A, Huang KW, Philson AC, Goldman M, Macosko EZ, McCarroll SA, Sabatini BL. 2017. Genetically Distinct Parallel Pathways in the Entopeduncular Nucleus for Limbic and Sensorimotor Output of the Basal Ganglia. Neuron 94:138–152 e5.

Wickens JR, Reynolds JN, Hyland BI. 2003. Neural mechanisms of reward-related motor learning. Curr Opin Neurobiol 13:685–690.

Williams H. 2004. Birdsong and singing behavior. Ann N Y Acad Sci 1016:1–30.

Williams H, Kilander K, Sotanski ML. 1993. Untutored song, reproductive success and song learning. Anim Behav 45:695–705.

Xiao L, Chattree G, Oscos FG, Cao M, Wanat MJ, Roberts TF. 2018. A Basal Ganglia Circuit Sufficient to Guide Birdsong Learning. Neuron 98:208–221 e5.

Zann RA. 1996. The zebra finch: a synthesis of field and laboratory studies. Oxford University Press.

